# Ancient coexistence of norepinephrine, tyramine, and octopamine signaling in bilaterians

**DOI:** 10.1101/063743

**Authors:** Philipp Bauknecht, Gáspár Jékely

## Abstract

Norepinephrine/noradrenaline is a neurotransmitter implicated in arousal and other aspects of vertebrate behavior and physiology. In invertebrates, adrenergic signaling is considered absent and analogous functions are performed by the biogenic amines octopamine and its precursor tyramine. These chemically similar transmitters signal by related families of GPCR in vertebrates and invertebrates, suggesting that octopamine/tyramine are the invertebrate equivalents of vertebrate norepinephrine. However, the evolutionary relationships and origin of these transmitter systems remain unclear. Using phylogenetic analysis and receptor pharmacology, here we establish that norepinephrine, octopamine, and tyramine receptors coexist in some marine invertebrates. In the protostomes *Platynereis dumerilii* (an annelid) and *Priapulus caudatus* (a priapulid) we identified and pharmacologically characterized adrenergic α1 and α2 receptors that coexist with octopamine α, octopamine β, tyramine type 1, and tyramine 2 receptors. These receptors represent the first examples of adrenergic receptors in protostomes. In the deuterostome *Saccoglossus kowalewskii* (a hemichordate), we identified and characterized octopamine α, octopamine β, tyramine type 1, and tyramine 2 receptors, representing the first example of these receptors in deuterostomes. *S. kowalewskii* also has adrenergic α1 and α2 receptors, indicating that all three signaling systems coexist in this animal. In phylogenetic analysis, we also identified adrenergic and tyramine receptor orthologs in xenacoelomorphs. Our results clarify the history of monoamine signaling in bilaterians. Since all six receptor families (two each for octopamine and tyramine and three for norepinephrine) can be found in representatives of the two major clades of Bilateria, the protostomes and the deuterostomes, all six receptors coexisted in the protostome-deuterostome last common ancestor. Adrenergic receptors were lost from most insects and nematodes and tyramine and octopamine receptors were lost from most deuterostomes. This complex scenario of differential losses cautions that octopamine signaling in protostomes is not a good model for adrenergic signaling in deuterostomes, and that the studies of marine animals where all three transmitter systems coexist will be needed for a better understanding of the origin and ancestral functions of these transmitters.

## Background

Norepinephrine is a major neurotransmitter in vertebrates with a variety of functions including roles in promoting wakefulness and arousal [1], regulating aggression [2], and autonomic functions such a heart beat [3]. Signaling by the monoamine octopamine in protostome invertebrates is often considered equivalent to vertebrate adrenergic signaling [4] with analogous roles in promoting aggression and wakefulness in flies [5, 6], or the regulation of heart rate in annelids and arthropods [7, 8]. Octopamine is synthesized from tyramine (Figure 1A) which itself also acts as a neurotransmitter or neuromodulator in arthropods and nematodes [4, 9-15]. Octopamine and norepinephrine are chemically similar, are synthesized by homologous enzymes [16, 17], and signal by similar but not orthologous G-protein coupled receptors (GPCRs) [4, 18].

Tyramine also signals by non-orthologous receptors in invertebrates and vertebrates. In insects and nematodes, tyramine signals by a GPCR that is related to octopamine receptors [12, 19]. In vertebrates, tyramine is only present at low levels and signals by the trace-amine receptors, a vertebrate-specific GPCR family only distantly related to the invertebrate tyramine receptors [20, 21]. Given these differences, the precise evolutionary relationships of these monoamine signaling systems are unclear.

The evolution of neurotransmitter systems has been analyzed by studying the distribution of monoamines or biosynthetic enzymes in different organisms [22]. This approach has limitations, however, because some of the biosynthetic enzymes are not specific to one substrate [16] and because trace amounts of several monoamines are found across many organisms, even if specific receptors are often absent [22]. For example, even if invertebrates can synthesize trace amounts of norepinephrine, these are not considered to be active neuronal signaling molecules, since the respective receptors are lacking. Consequently, the presence of specific monoamine receptors is the best indicator that a particular monoamine is used in neuronal signaling [11, 23].

To clarify the evolutionary history of adrenergic, octopamine, and tyramine signaling in animals, we decided to undertake a comparative phylogenetic and pharmacological study of these receptor families in bilaterians. Bilaterians, animals with bilateral symmetry, are comprised of protostomes, deuterostomes, and xenacoelomorphs [24]. Deuterostomes include chordates and ambulacrarians (hemichordates and echinoderms), and protostomes are formed by the clades Ecdysozoa, Lophotrochozoa (Spiralia), and Chaetognatha. Ecdysozoa includes arthropods, nematodes, priapulids and other phyla. Lophotrochozoa include annelids, mollusks, and other, mostly marine groups. Xenacoelomorps, a group including acoel flatworms, nemertodermatids, and *Xenoturbella*, have been proposed to belong to the deuterostomes, or represent a sister group to all remaining bilaterians [25-27]. Here we establish the orthology relationships of adrenergic, octopamine, and tyramine receptors across bilaterians. We find that six receptor families originated at the base of the bilaterian tree. We then pharmacologically characterize adrenergic receptors from an annelid and a priapulid, and octopamine and tyramine receptors from an annelid and a hemichordate. The broad phylogenetic sampling and comparative pharmacology paints a richer picture of the evolution of these receptors, characterized by ancestral coexistence and multiple independent losses.

## Results

Using database searches, sequence-similarity-based clustering, and phylogenetic analysis, we reconstructed the phylogeny of α1, α2, and β adrenergic, octopamine α, octopamine β, and tyramine type-1 and type-2 receptors. Each family formed well-resolved clusters in a sequence-similarity-based clustering analysis and well-supported clades in molecular phylogenetic analysis (Figure 1B, C and Additional file 1).

We identified several invertebrate GPCR sequences that were similar to vertebrate adrenergic α1 and α2 receptors (Figure 1B, C). An adrenergic α1 receptor ortholog is present in the sea urchin *Strongylocentrotus purpuratus.* Adrenergic α1 and α2 receptors were both present in *Saccoglossus kowalewskii*, a hemichordate deuterostome (Figure 1B, C and Additional files 1-3), as previously reported [28]. We also identified adrenergic α1 and α2 receptor orthologs in annelids and mollusks (members of the Lophotrochozoa), including *Aplysia californica*, and in the priapulid worm *Priapulus caudatus* (member of the Ecdysozoa)(Figure 1B, C and Additional files 1-3). Adrenergic α receptors are also present in a few arthropods, including the crustacean *Daphnia pulex* and the moth *Chilo suppressalis* (the *Chilo* α2 receptor was first described as an octopamine receptor [29]), but are absent from most other insects (Additional files 1-3). Adrenergic α2 receptors are also present in xenacoelomorphs, in *Xenoturbella bocki* and the nemertodermatid *Meara stichopi. M. stichopi* also has two adrenergic α1 receptor orthologs (Figure 1C and Additional file 1-3).

**Figure 1.**
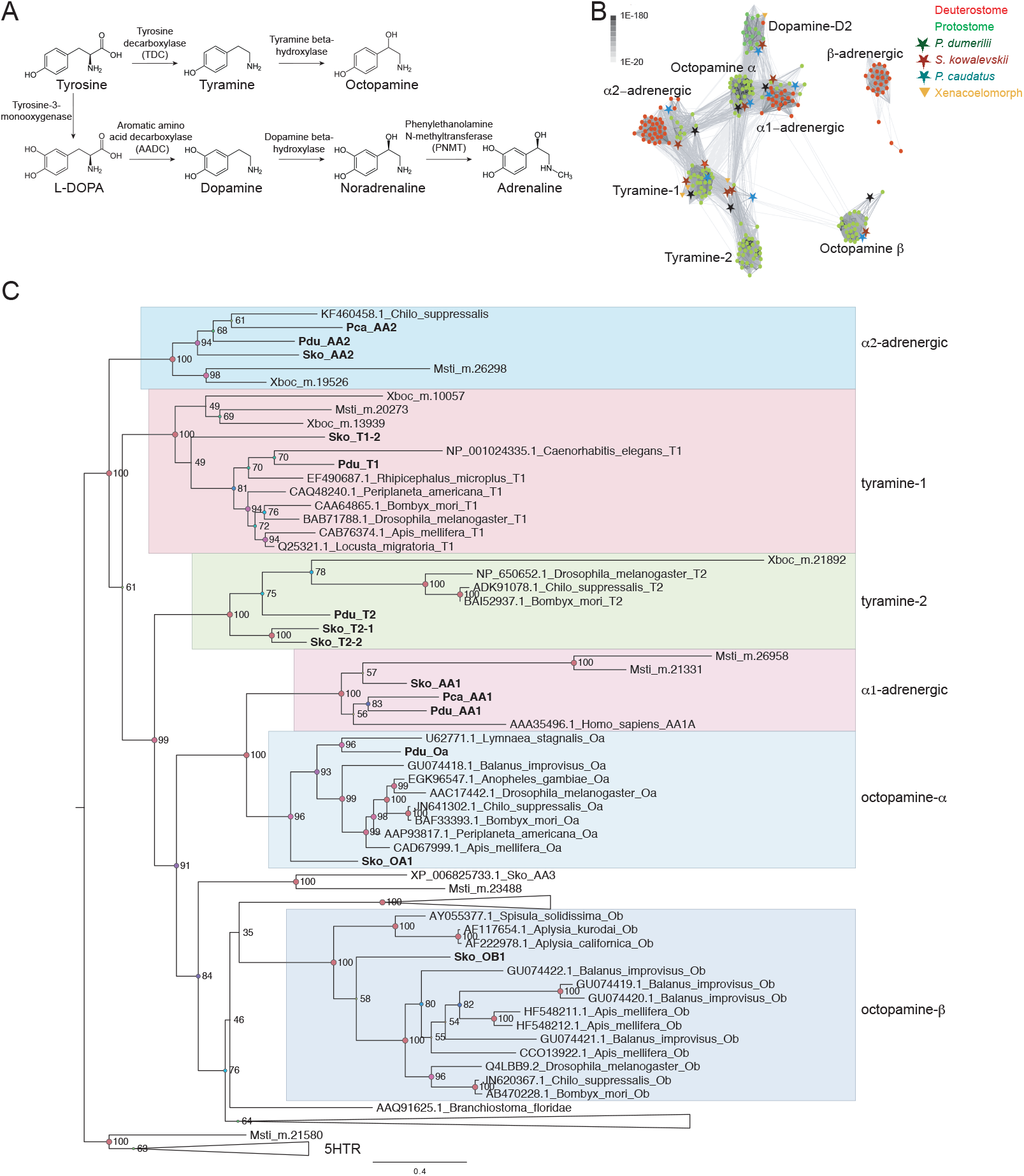
Biosynthesis of monoamines and phylogeny of adrenergic, tyramine, and octopamine GPCR sequences. (A) Biosynthesis of tyramine, octopamine, norepinephrine, and epinephrine from tyrosine. The enzymes catalyzing the reaction steps are indicated. (B) Sequence-similarity-based cluster map of bilaterian octopamine, tyramine, and adrenergic GPCRs. Nodes correspond to individual GPCRs and are colored based on taxonomy. Edges correspond to BLAST connections of P value >1e-70. (C) Simplified phylogenetic tree of bilaterian adrenergic, tyramine, and octopamine GPCR sequences. The tree is rooted on 5HT receptors. Abbreviations: Pdu, *P. dumerilii;* Pca, *P. caudatus;* Sko, *S. kowalevskii*; Msti, *M. stichopi;* Xboc, *X. bocki.*

The identification of adrenergic α1, and α2 receptor orthologs in ambulacrarians, lophotrochozoans, ecdysozoans, and xenacoelomorphs indicates that both families were present in the bilaterian last common ancestor.

Adrenergic β receptors are found in chordates, including urochordates and cephalochordates. In addition, we identified an adrenergic β receptor ortholog in the xenacoelomorph *M. stichopi* (Additional file 4). If xenacoelomorphs are sister to all remaining bilaterians, then this receptor family also originated at the base of Bilateria and was lost from all protostomes. To characterize the ligand specificities of these putative invertebrate adrenergic receptors, we cloned them from *S. kowalewskii, P. caudatus*, and the marine annelid *Platynereis dumerilii.* We performed *in vitro* GPCR activation experiments using a Ca^2+^-mobilization assay [30, 31]. We found that norepinephrine and epinephrine activated both the adrenergic α1 and α2 receptors from all three species with EC_50_ values in the high nanomolar range or lower. In contrast, tyramine, octopamine, and dopamine were either inactive or only activated the receptors at approximately two orders of magnitude higher concentrations (Figure 2, Table 1). These phylogenetic and pharmacological results collectively establish these invertebrate receptors as bona fide adrenergic α receptors.

**Figure 2.**
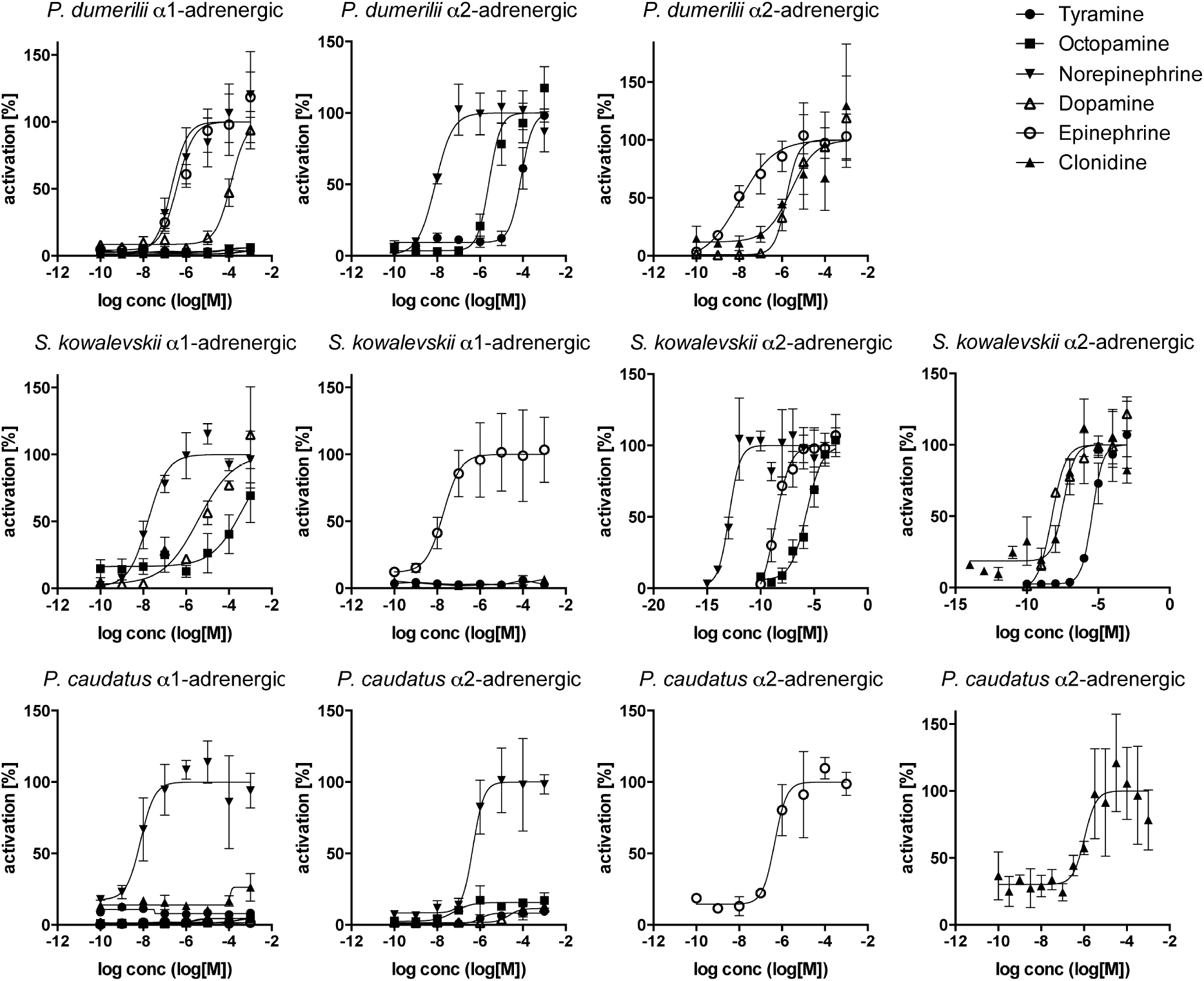
Dose-Response curves of adrenergic GPCRs from *P. dumerilii, P. caudatus*, and *S. kowalevskii* treated with varying concentrations of ligand. Data, representing luminescence units relative to the maximum of the fitted dose-response curves, are shown as mean ± SEM (n = 3). EC50 values and significance values are listed in Table 1.

**Table 1.** EC_50_ (M)/IC_5_o (M) values of all tested GPCRs with the indicated ligands or inhibitors. The most effective natural ligand for each receptor is shown in bold. 95% confidence intervals for the EC_50_ (M)/IC_50_ (M) values are given in every second line. The lowest EC_50_ value for each receptor was compared to the next lowest one using the extra sum-of-squares F test. ***, p<0.0001; *, p<0.05; n.s., not significant. Significance values are shown for the compared pairs.

| EC<sub>50</sub> (M)/IC<sub>50</sub> (M) | Tyramine | Octopamine | Clonidine | Norepinephrine | Dopamine | Epinephrine | Yohimbine | Mianserin |
| --- | --- | --- | --- | --- | --- | --- | --- | --- |
| *P. dumerilii* α1-adrenergic | inactive | inactive | inactive | **2.1E-07** *** | 1.2E-04 *** | 3.7E-07 n.s. | 4.4E-06 | 3.7E-06 |
| 95% CI |  |  |  | 1.0E-007 to 4.2E-007 | 2.7E-005 to 0.00056 | 1.3E-007 to 1.1E-006 | 2.3E-006 to 8.2E-006 | 1.9E-006 to 7.2E-006 |
| *P. dumerilii* α2-adrenergic | 8.4E-05 | 2.7E-06 *** | 2.6E-06 | **8.2E-09** *** | 1.6E-06 | 1.1E-08 n.s. | 5.7E-06 | 2.5E-05 |
| 95% CI | 2.8E-005 to 0.00024 | 6.683E-007 to 1.0E-005 | 2.4E-007 to 2.7E-005 | 5.7E-009 to 1.1E-008 | 8.3E-007 to 3.2E-006 | 5.0E-009 to 2.2E-008 | 3.5E-006 to 9.1E-006 | 1.2E-005 to 5.1E-005 |
| *S. kowalevskii* α1-adrenergic | inactive | inactive | inactive | **1.7E-08** *** | 3.8E-06 *** | 1.9E-08 n.s. | 1.3E-05 | 4.5E-06 |
| 95% CI |  |  |  | 1.0E-008 to 2.7E-008 | 1.9E-007 to 7.4E-005 | 9.0E-009 to 4.1E-008 | 7.6E-006 to 2.2E-005 | 1.6E-006 to 1.1E-005 |
| *S. kowalevskii* α2-adrenergic | 3.7E-06 | 1.9E-06 | 3.6E-08 | **1.2E-13** *** | 5.6E-09 | 2.3E-09 *** | 3.3E-07 | inactive |
| 95% CI | 2.0E-006 to 6.8E-006 | 2.5E-007 to 1.4E-005 | 6.7E-009 to 1.9E-007 | 6.7E-014 to 1.9E-013 | 3.3E-009 to 9.4E-009 | 1.1E-009 to 4.6E-009 | 2.6E-007 to 4.0E-007 |  |
| *P. caudatus* α1-adrenergic | inactive | inactive | inactive | **7.5E-09** | inactive | inactive | inactive | inactive |
| 95% CI |  |  |  | 4.0E-009 to 1.3E-008 |  |  |  |  |
| *P. caudatus* α2-adrenergic | inactive | inactive | 1.1E-06 *<br>p=0.021 | 4.7E-07 * | inactive | **4.5E-07** n.s. | inactive | 9.8E-07 |
| 95% CI |  |  | 4.5E-007 to 2.4E-006 | 1.7E-007 to 1.2E-006 |  | 1.8E-007 to 1.0E-006 |  | 4.3E-007 to 2.2E-006 |
| *P. dumerilii* Tyramine-1 | **1.1E-08** *** | 2.7E-06 *** | 2.1E-06 | 1.7E-05 | 7.8E-06 | 3.1E-05 | 2.1E-06 | 4.7E-05 |
| 95% CI | 7.6E-009 to 1.6E-008 | 1.1E-006 to 6.1E-006 | 1.0E-006 to 4.1E-006 | 1.0E-005 to 2.8E-005 | 1.5E-006 to 3.9E-005 | 9.8E-006 to 9.9E-005 | 7.0E-007 to 6.0E-006 | 1.7E-005 to 0.00012 |
| *P. dumerilii* Tyramine-2 | **7.0E-09** *** | 7.8E-07 *** | 5.3E-06 | 1.1E-04 | 3.9E-06 | 4.8E-05 | 5.4E-05 | 6.4E-06 |
| 95% CI | 3.0E-009 to 1.6E-008 | 3.8E-007 to 1.5E-006 | 2.1E-006 to 1.3E-005 | 2.9E-005 to 0.00038 | 2.1E-006 to 7.0E-006 | 8.6E-006 to 0.00026 | 3.6E-005 to 7.9E-005 | 3.9E-006 to 1.0E-005 |
| *S. kowalevskii* Tyramine-1 | **8.6E-05** n.s. | inactive | 2.9E-04 n.s. | inactive | 0.57 | inactive | 1.7E-06 | 1.7E-05 |
| 95% CI | 2.8E-005 to 0.00025 |  | 0.00013 to 0.00065 |  | very wide | 2.1E-006 to 0.00017 | 7.1E-007 to 3.9E-006 | 7.7E-006 to 3.7E-005 |
| *S. kowalevskii* Tyramine-2A | **1.0E-09** *** | 8.6E-08 *** | 1.4E-06 | inactive | 7.2E-08 | inactive | inactive | 1.6E-04 |
| 95% CI | 6.6E-010 to 1.5E-009 | 4.0E-008 to 1.8E-007 | 7.4E-007 to 2.6E-006 |  | 1.4E-008 to 3.5E-007 |  |  | 5.4E-008 to 0.47 |
| *S. kowalevskii* Tyramine-2B | **5.9E-09** *** | 1.6E-06 *** | 1.6E-05 | 1.2E-04 | 1.4E-06 | 2.8E-05 | 2.1E-05 | 1.9E-05 |
| 95% CI | 2.4E-009 to 1.4E-008 | 6.5E-007 to 3.7E-006 | 6.2E-006 to 4.0E-005 | 3.6E-005 to 0.00036 | 9.0E-007 to 2.2E-006 | 5.1E-006 to 0.00015 | 1.1E-005 to 3.6E-005 | 1.1E-005 to 3.0E-005 |
| *P. dumerilii* Octopamine α | 1.3E-05 | **2.6E-07** * | 1.4E-07 n.s. | 3.5E-06 *<br>p=0.003 | inactive | 8.8E-06 | 9.0E-09 | 1.6E-06 |
| 95% CI | 4.2E-006 to 4.1E-005 | 8.4E-008 to 7.7E-007 | 6.7E-008 to 3.0E-007 | 1.8E-006 to 6.7E-006 |  | 2.5E-006 to 3.0E-005 | 4.1E-009 to 1.9E-008 | 9.7E-007 to 2.6E-006 |
| *S. kowalevskii* Octopamine α | 1.7E-05 | **6.9E-07** * | 1.6E-07 *<br>p=0.048 | 5.3E-05 | 2.6E-04 | 1.8E-05 | 7.8E-06 | 2.2E-05 |
| 95% CI | 3.0E-006 to 9.5E-005 | 1.8E-007 to 2.4E-006 | 7.6E-008 to 3.5E-007 | 1.5E-005 to 0.00018 | 3.4E-006 to 0.02 | 7.1E-006 to 4.7E-005 | 3.1E-006 to 1.8E-005 | 1.2E-005 to 3.6E-005 |
| *S. kowalevskii* Octopamine β | inactive | **6.4E-08** *** | inactive | 3.5E-06 *** | inactive | inactive | 1.6E-04 | 6.4E-06 |
| 95% CI |  | 4.0E-008 to 1.0E-007 |  | 1.4E-006 to 8.1E-006 |  |  | 1.0E-005 to 0.0023 | 3.1E-006 to 1.3E-005 |

To investigate if adrenergic signaling coexists with octopamine and tyramine signaling in protostomes, we searched for octopamine and tyramine receptors in *P. dumerilii* and *P. caudatus.* In phylogenetic and clustering analyses, we identified orthologs for tyramine type 1 and type 2 and octopamine α and β receptors in both species (Figure 1B, C and Additional files 5-8). We performed activation assays with the *P. dumerilii* receptors. The tyramine type 1 and type 2 receptors orthologs were preferentially activated by tyramine with EC_50_ values in the nanomolar range (Figure 3, Table 1). The *P. dumerilii* octopamine α receptor was activated by octopamine at a lower concentration than by tyramine and dopamine (Figure 4, Table 1). The *P. dumerilii* octopamine β receptor was not active in our assay. These results show that specific receptor systems for norepinephrine, octopamine, and tyramine coexist in *P. dumerilii* and very likely also *P. caudatus.*

**Figure 3.**
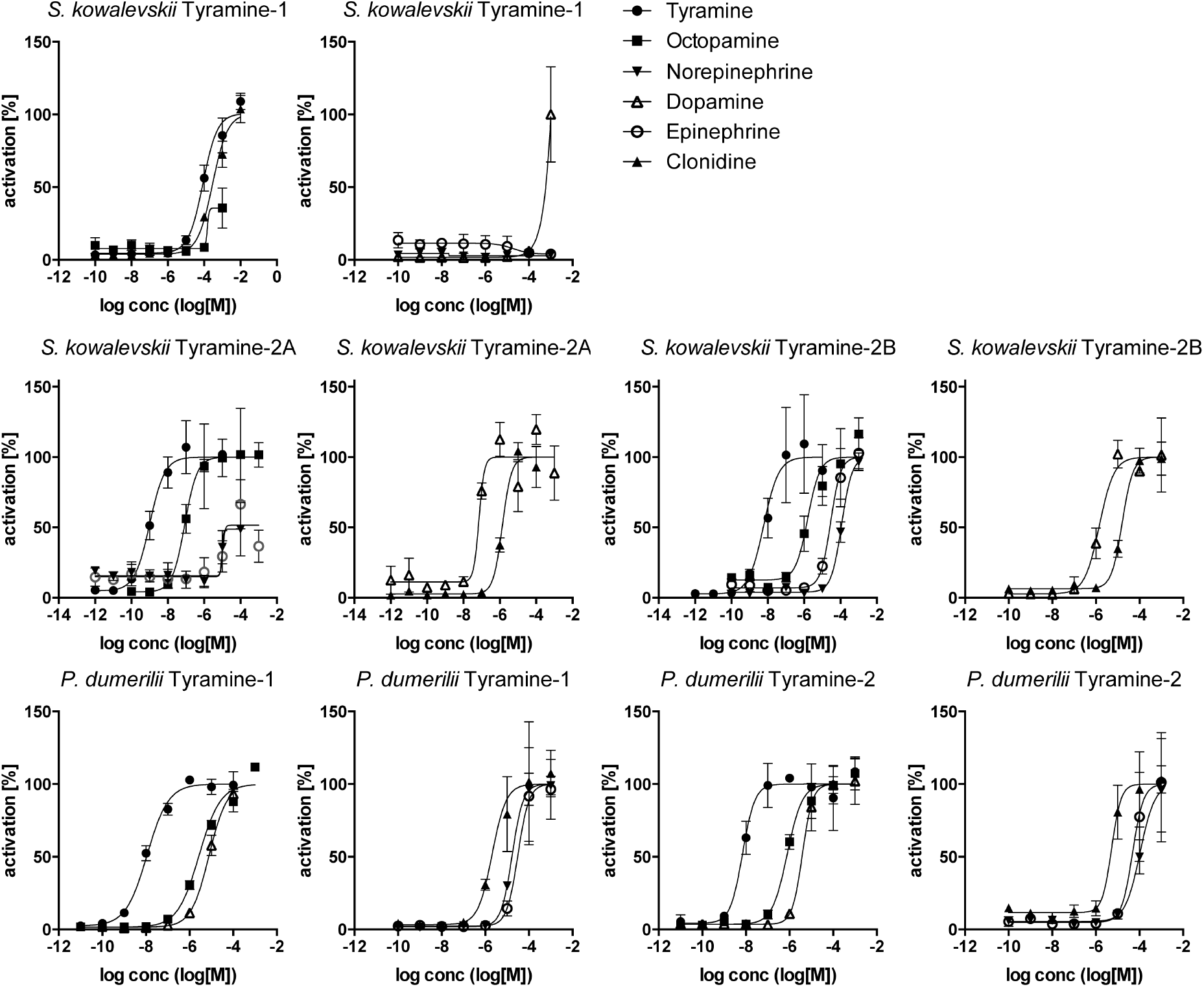
Dose-Response curves of tyramine GPCRs from *P. dumerilii* and *S. kowalevskii* treated with varying concentrations of ligand. Data, representing luminescence units relative to the maximum of the fitted dose-response curves, are shown as mean ± SEM (n = 3). EC50 values and significance values are listed in Table 1.

When did tyramine and octopamine signaling originate? To answer this, we surveyed available genome sequences for tyramine and octopamine receptors. As expected, we identified several receptors across the protostomes, including ecdysozoans and lophotrochozoans (Additional files 5-8). We also identified tyramine, but not octopamine, receptors in xenacoelomorphs. However, chordate genomes lacked orthologs of these receptors. Strikingly, we identified tyramine type 1 and 2 and octopamine α and β receptor orthologs in the genome of the hemichordate *S. kowalewskii* (Figure 1B, C, Additional files 5-8). In phylogenetic analyses, we recovered at least one *S. kowalewskii* sequence in each of the four receptor clades (one octopamine α, one octopamine β, two tyramine type 1, and two tyramine type 2 receptors), establishing these sequences as deuterostome orthologs of these predominantly protostome GPCR families (Additional files 5-8).

We cloned the candidate *S. kowalewskii* tyramine and octopamine receptors and performed ligand activation experiments. The *S. kowalewskii* type 2 receptors were preferentially activated by tyramine in the nanomolar range. The type 1 receptor was only activated at higher ligand concentrations. The octopamine α and β receptors were preferentially activated by octopamine in the nanomolar range (Figures 3 and 4, Table 1). These data show that octopamine and tyramine signaling also coexists with adrenergic signaling in this deuterostome, as in *P. dumerilii* and *P. caudatus.* The presence of tyramine signaling in *S. kowalewskii* is also supported by the phylogenetic distribution of tyrosine decarboxylase, a specific enzyme for tyramine synthesis [32]. Tyrosine decarboxylase is present in protostomes and *S. kowalewskii* but is absent from other deuterostomes (Additional file 9). In mammals, aromatic amino acid decarboxylases are involved in synthesizing low amounts of tyramine [33].

**Figure 4.**
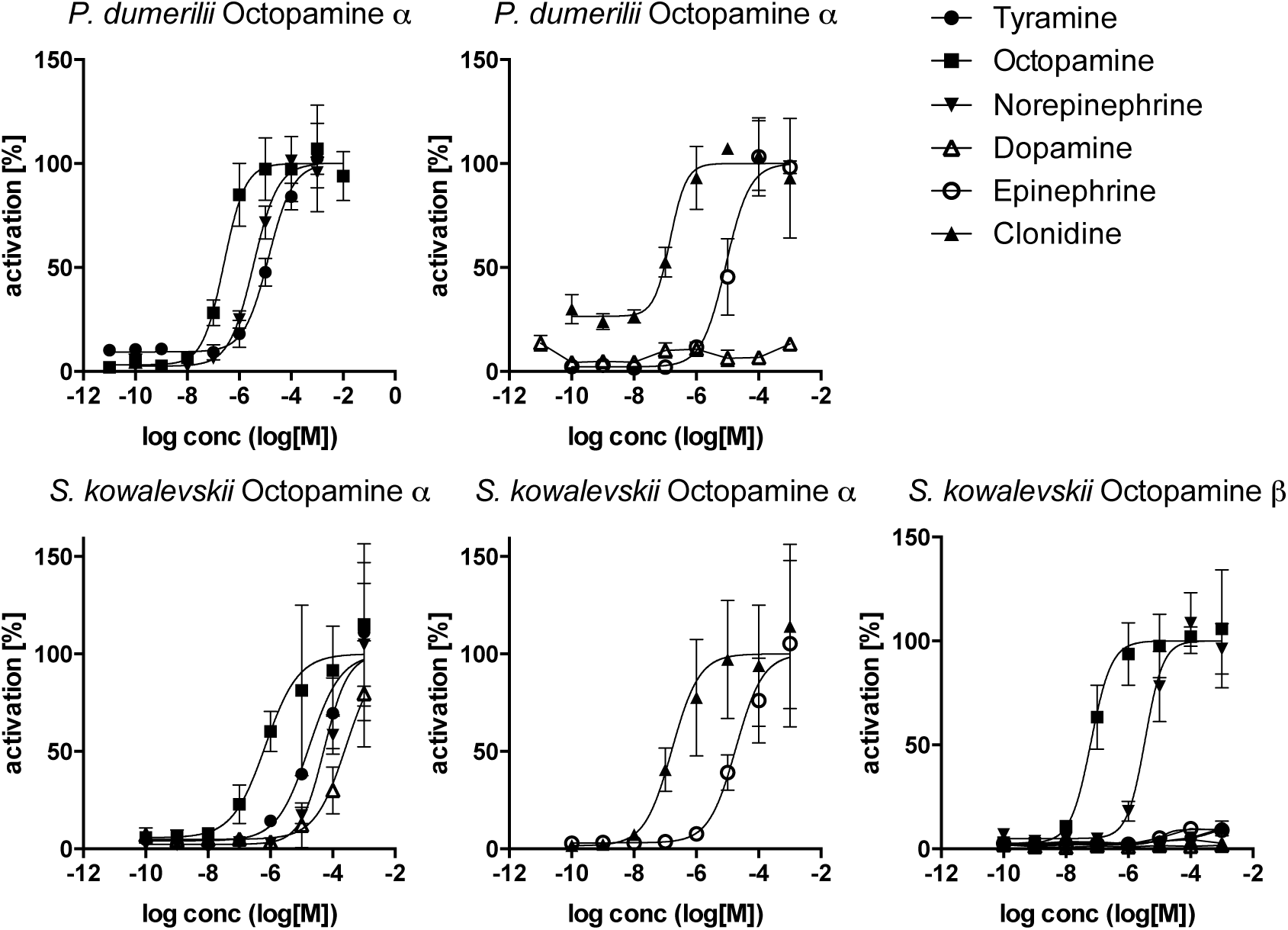
Dose-Response curves of octopamine GPCRs from *P. dumerilii* and *S. kowalevskii* treated with varying concentrations of ligand. Data, representing luminescence units relative to the maximum of the fitted dose-response curves, are shown as mean ± SEM (n = 3). EC_50_ values and significance values are listed in Table 1.

We also tested the α adrenergic agonist clonidine and the GPCR antagonists mianserin and yohimbine on several receptors from all three species. These chemicals did not show specificity for any of the receptor types, suggesting these chemicals may not be useful for studying individual biogenic amine receptors *in vivo* (Table 1. and Additional file 10).

## Discussion

The discovery of adrenergic signaling in some protostomes and xenacoelomorphs and octopamine and tyramine signaling in a deuterostome changes our view on the evolution of monoamine signaling in bilaterians (Figure 5). It is clear from the phylogenetic distribution of orthologous receptor systems that at least six families of octopamine, tyramine, and adrenergic receptors were present in the bilaterian last common ancestor. These include the adrenergic α1 and α2 receptors, the tyramine type 1 and type 2 receptors, and the octopamine α and β receptors. From the six ancestral families, the octopamine and tyramine receptors were lost from most deuterostomes, and the adrenergic receptors were lost from most ecdysozoans. Interestingly, the xenacoelomorph *M. stichopi* also has an adrenergic β receptor, representing the only ortholog outside chordates. Octopamine α receptors were likely lost from xenacoelomorphs, since the split of the six receptor families (four with well-resolved xenacoelomorph sequences) predated the divergence of the main lineages of bilaterians (Figure 1C).

**Figure 5.**
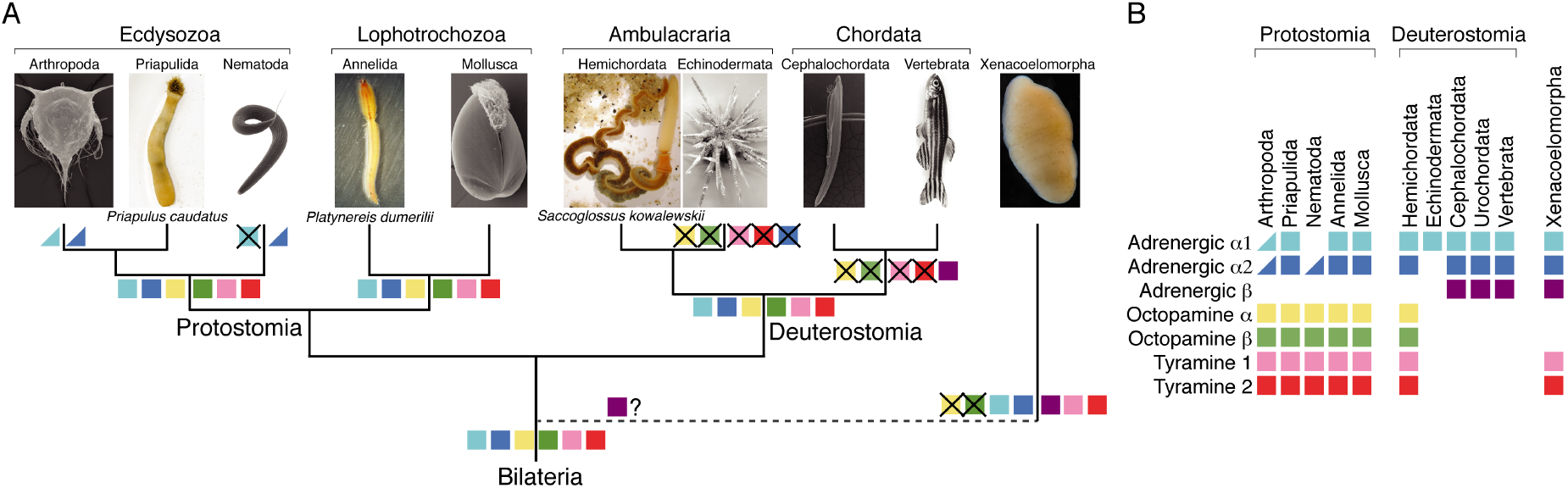
Evolution of adrenergic, octopamine, and tyramine signaling in bilaterians. (A) Phylogenetic tree of major clades of bilaterian animals with the presence/loss of specific GPCR families indicated. (B) Phyletic distribution of adrenergic, octopamine, and tyramine GPCR families across major bilaterian clades. Half squares mean losses in a large number of species in a phylum.

Although we performed the receptor activation assays in a heterologous system that may not mimic the *in vivo* situation very well, we find clear evidence of ligand preferences for each receptor. In general, there is a two orders of magnitude difference in the EC_50_ values between the best ligand and other related ligands for the same receptor measured under the same conditions. We consider these *in vitro* ligand preferences as indicative of the physiological ligands for these receptors. Furthermore, there is a high congruence between the *in vitro* ligand specificities and the phylogenetic placement of the different classes of receptors, further strengthening our receptor-type assignments. The most potent ligand of all six orthologous receptor families we analyzed is the same across protostomes and deuterostomes, indicating the evolutionary stability of ligand-receptor pairs, similar to the long-term stability of neuropeptide GPCR ligand-receptor pairs [34, 35].

Understanding the ancestral role of these signaling systems and why they may have been lost differentially in different animal groups will require functional studies in organisms where all three neurotransmitter systems coexist.

## Conclusions

We established the coexistence of adrenergic, octopaminergic, and tyraminergic signaling in the deuterostome *S. kowalewskii* and the protostomes *P. dumerilii* and *P. caudatus.* Signaling by norepinephrine in vertebrates has often been considered as equivalent to signaling by octopamine in invertebrates. Our results change this view and show that these signaling systems coexisted ancestrally and still coexist in some bilaterians. The extent of functional redundancy in species where all six receptor systems coexist will require experimental studies. It may be that some of these monoamines ancestrally had partially overlapping roles. In that case, following the loss of a receptor, functions associated with that ligand-receptor pair may have been taken over by another pair. However, regardless of such potential shifts in function, it is clear that octopamine signaling in invertebrates and adrenergic signaling in vertebrates is not equivalent or homologous from an evolutionary point of view. This has important implications for our interpretation of comparative studies of the function of these neurotransmitter systems and their neural circuits. Our study also contributes to the understanding of nervous system evolution in bilaterians by revealing extensive losses during the history of one of the major classes of neurotransmitter systems.

## Methods

### Gene identification and receptor cloning

*Platynereis* protein sequences were collected from a *Platynereis* mixed stages transcriptome assembly [36]. GPCR sequences from other species were downloaded from NCBI. GPCRs were cloned into pcDNA3.1(+) (Thermo Fisher Scientific. Waltham. USA) as described before [31]. Forward primers consisted of a spacer (ACAATA) followed by a BamHI or EcoRI restriction site, the Kozak consensus sequence (CGCCACC), the start codon (ATG) and a sequence corresponding to the target sequence. Reverse primers consisted of a spacer (ACAATA), a NotI restriction site, a STOP codon, and reverse complementary sequence to the target sequence. Primers were designed to end with a C or G with 72°C melting temperature. PCR was performed using Phusion polymerase (New England Biolabs GmbH, Frankfurt, Germany). The sequences of all *Platynereis* GPCRs tested here were deposited in GenBank (accession numbers: α1-adrenergic receptor. KX372342; α2-adrenergic receptor, KX372343 Tyramine-1 receptor, KP293998; Tyramine-2 receptor, KU715093; Octopamine α receptor, KU530199; Octopamine β receptor, KU886229). Tyramine receptor 1 has been previously published [31] as Pdu orphan GPCR 48. The GenBank accession numbers of the *S. kowalevskii* and *P. caudatus* sequences tested are: *S. kowalevskii* α1-adrenergic, ALR88680; S. kovalewskii α2-adrenergic, XP_002734932; *P. caudatus* α1-adrenergic XP_014662992; *P. caudatus* α2-adrenergic, XP_014681069; *S. kovalewskii* Tyramine-1, XP_002742354; *S. kovalewskii* Tyramine-2A, XP_002734062; S. kovalewskii Tyramine-2B, XP_006812999; *S. kowalevskii* Octopamine α, XP_006823182; *S. kowalevskii* Octopamine β, XP_002733926.

### Cell culture and receptor deorphanization

Cell culture assays were done as described before [31]. Briefly, CHO-K1 cells were kept in Ham's F12 Nut Mix medium (Thermo Fisher Scientific, Waltham, USA) with 10 % fetal bovine serum and penicillin-streptomycin (PenStrep, Invitrogen). Cells were seeded in 96-well plates (Thermo Fisher Scientific, Waltham, USA) at approximately 10,000 cells/well. After 1 day, cells were transfected with plasmids encoding a GPCR, the promiscuous Gα-16 protein [37], and a reporter construct GFP-apoaequorin [38] (60 ng each) using 0.375 μl of the transfection reagent TurboFect (Thermo Fisher Scientific. Waltham. USA). After two days of expression, the medium was removed and replaced with Hank's Balanced Salt Solution (HBSS) supplemented with 1.8 mM Ca^2^+, 10 mM glucose, and 1 mM coelenterazine h (Promega, Madison, USA). After incubation at 37°C for 2 hours, cells were tested by adding synthetic monoamines (Sigma, St. Louis, USA) in HBSS supplemented with 1.8 mM Ca^2^+ and 10 mM glucose. Solutions containing norepinephrine, epinephrine or dopamine were supplemented with 100 μM ascorbic acid to prevent oxidation. Luminescence was recorded for 45 seconds in a plate reader (BioTek Synergy Mx or Synergy H4, BioTek, Winooski, USA). For inhibitor testing, the cells were incubated with yohimbine or mianserin (Sigma, St. Louis, USA) for 1 hour. Then, synthetic monoamines were added to yield in each case the smallest final concentration expected to elicit the maximal response in the absence of inhibitor and luminescence was recorded for 45 seconds. Data were integrated over the 45-second measurement period. Data for dose-response curves were recorded in triplicate for each concentration. Dose-response curves were fitted with a four-parameter curve using Prism 6 (GraphPad, La Jolla. USA). The curves were normalized to the calculated upper plateau values (100% activation). The different EC50 values for each receptor were compared with the extra sum-of-squares F test in a pairwise manner using Prism 6.

### Bioinformatics

Protein sequences were downloaded from the NCBI. Redundant sequences were removed from the collection using CD-HIT [39] with an identity cutoff of 70%. Sequence cluster maps were created with CLANS2 [40] using the BLOSUM62 matrix and a P-value cutoff of 1e-70. For phylogenetic trees, protein sequences were aligned with MUSCLE [41]. Alignments were trimmed with TrimAI [42] in "Automated 1" mode. The best amino acid substitution model was selected using ProtTest 3 [43]. Maximum likelihood trees were calculated with RAxML [44] using the CIPRES Science Gateway [45] or with IQ-TREE and automatic model selection (http://www.iqtree.org/). Bootstrap analysis in RAxML was done and automatically stopped [46] when the Majority Rule Criterion (autoMRE) was met. The resulting trees were visualized with FigTree (http://tree.bio.ed.ac.uk/software/figtree/). The identifiers of deorphanized adrenergic, octopamine, and tyramine receptors [12, 29, 47-59] were tagged with _AA1, AA2, _Oa, _Ob, _T1, or _T2. The trees were rooted on 5HT receptors.

### Funding

The research leading to these results received funding from the European Research Council under the European Union's Seventh Framework Programme (FP7/2007-2013)/ European Research Council Grant Agreement 260821. PB is supported by the International Max Planck Research School (IMPRS) ''From Molecules to Organisms.''

The funding bodies had no role in the design of the study and collection, analysis, and interpretation of data, and in writing the manuscript.

## Acknowledgements

We thank John Gerhart for Saccoglossus DNA and the image of Saccoglossus. We thank Mattias Hogvall for Priapulus DNA and the image of Priapulus, and Anne-C. Zakrzewski for the *Xenoturbella* image. We also thank Elizabeth Williams for comments on the manuscript.

## Additional files

**Additional file 1.**
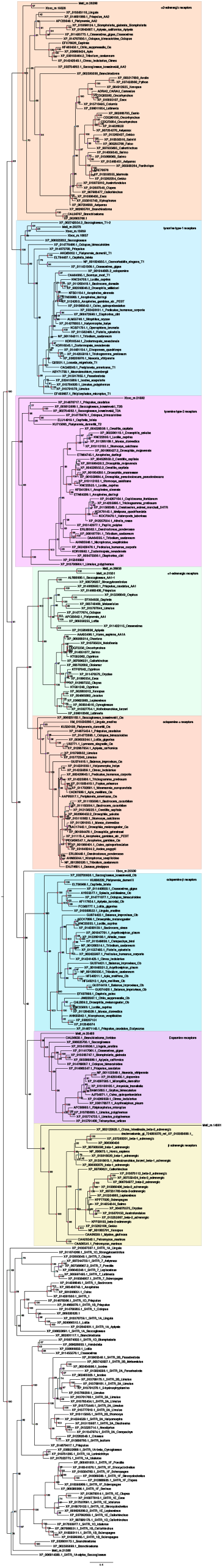
Maximum likelihood tree of adrenergic, octopamine, and tyramine receptors. Bootstrap support values are shown. This tree containing all investigated GPCRs. The tree was rooted on 5HT receptor sequences. Subtrees are shown in Additional files 2-8.

**Additional file 2.**
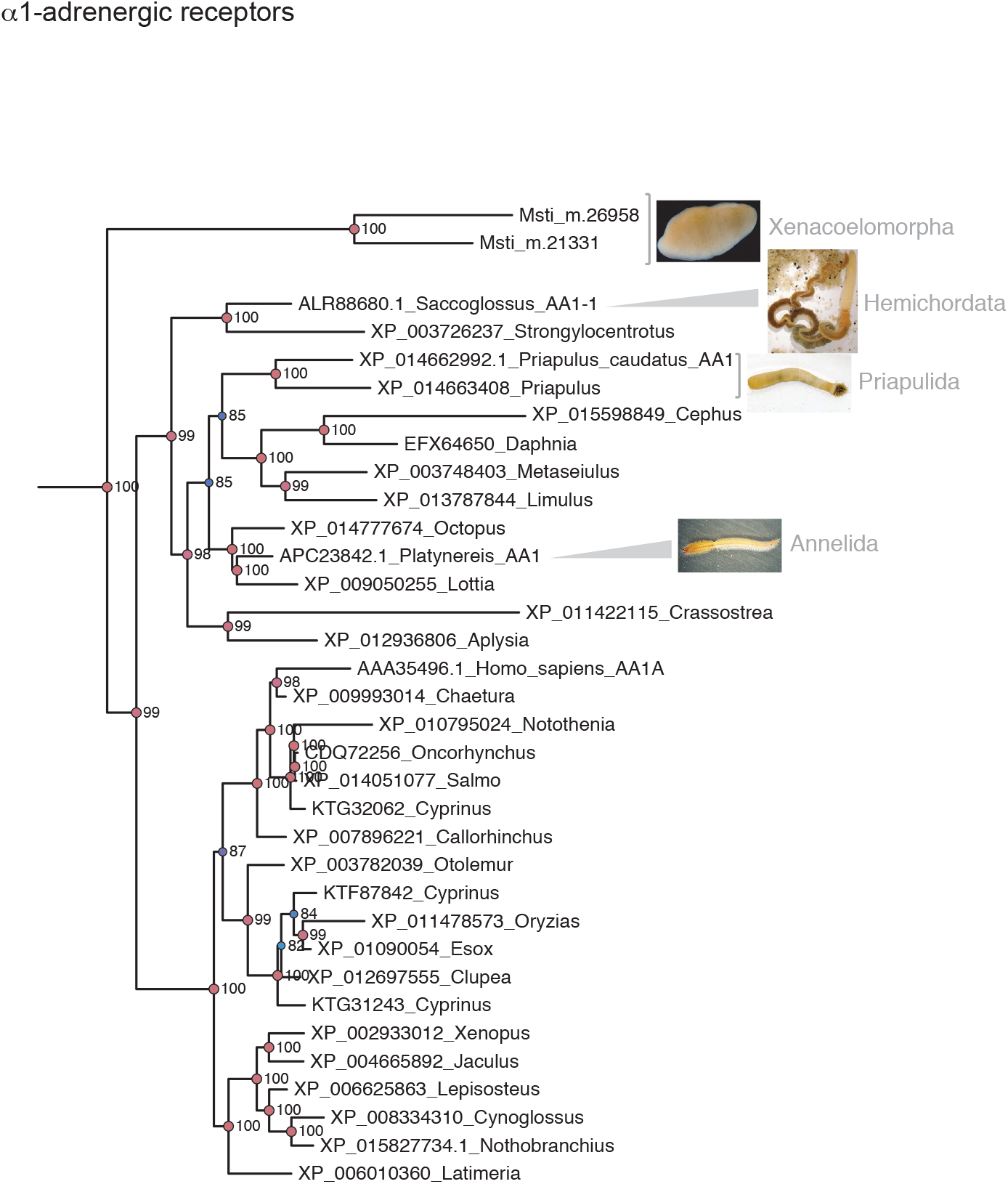
Maximum likelihood tree of α1-adrenergic receptors. Bootstrap support values are shown for selected nodes. This tree is part of a larger tree containing all investigated GPCRs.

**Additional file 3.**
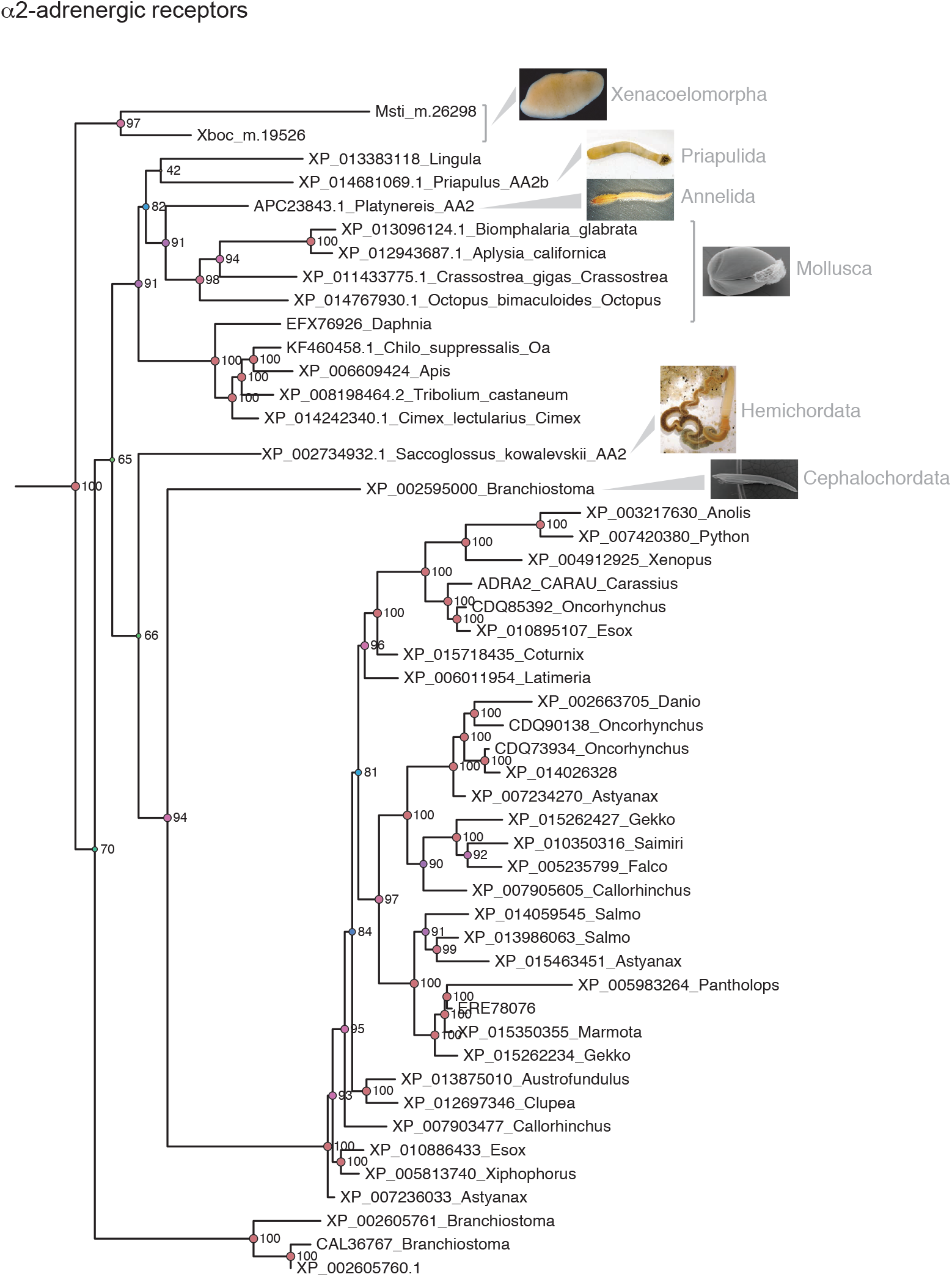
Maximum likelihood tree of a2-adrenergic receptors. Bootstrap support values are shown for selected nodes. This tree is part of a larger tree containing all investigated GPCRs.

**Additional file 4.**
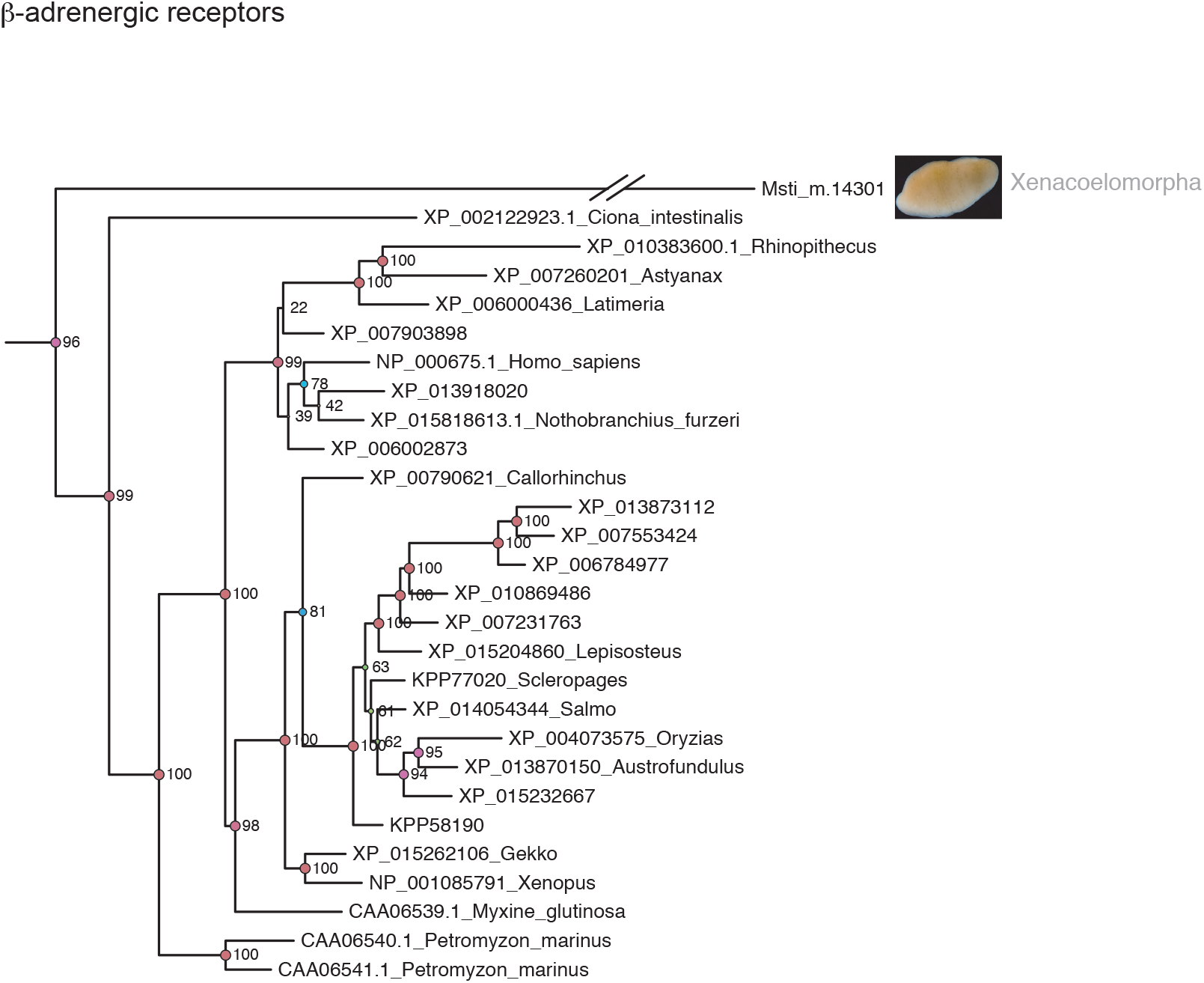
Maximum likelihood tree of β-adrenergic receptors. Bootstrap support values are shown for some nodes of interest. This tree is part of a larger tree containing all investigated GPCRs.

**Additional file 5.**
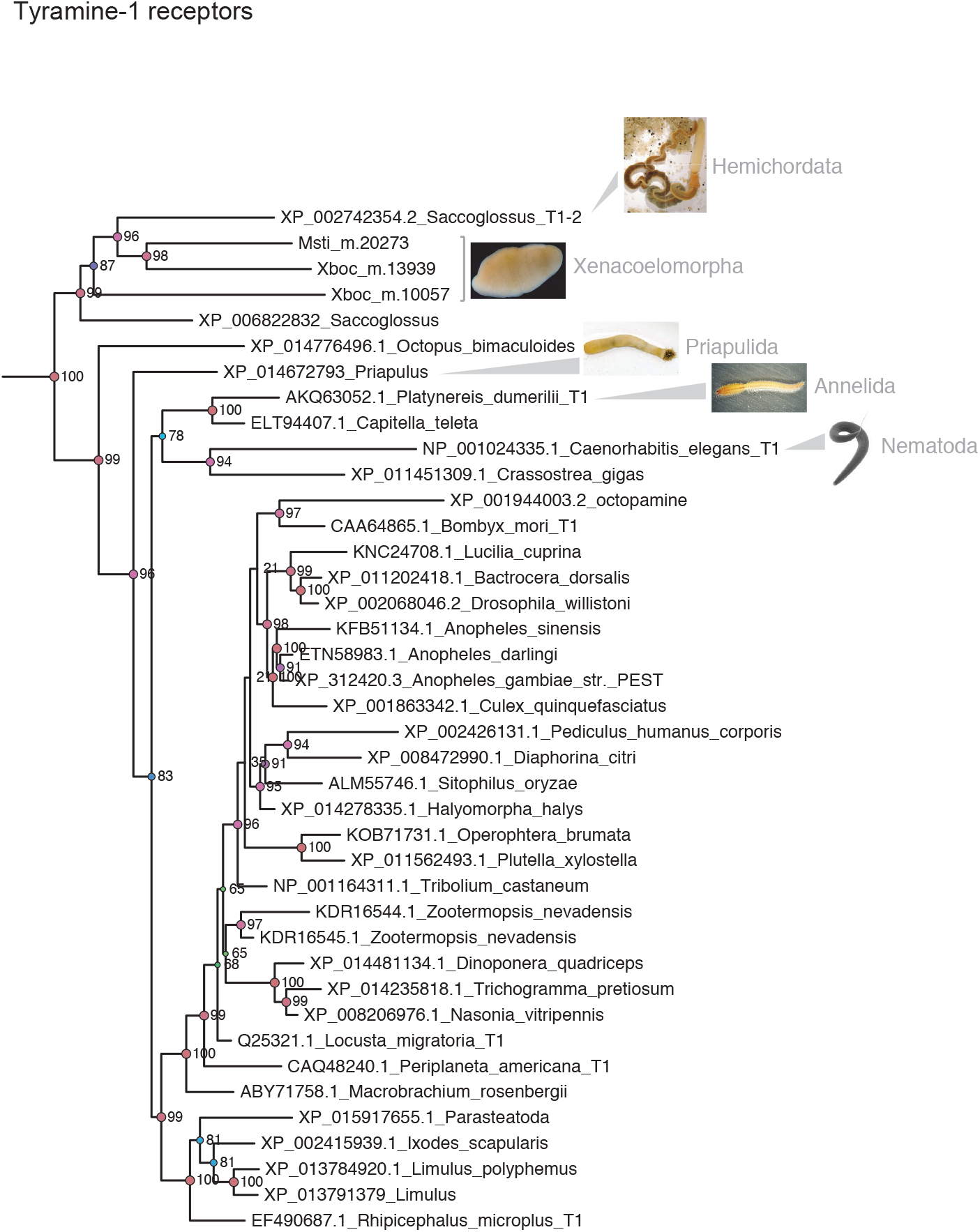
Maximum likelihood tree of Tyramine-1 receptors. Bootstrap support values are shown for selected nodes. This tree is part of a larger tree containing all investigated GPCRs. The identifiers of deorphanized tyramine receptors were tagged with _T1.

**Additional file 6.**
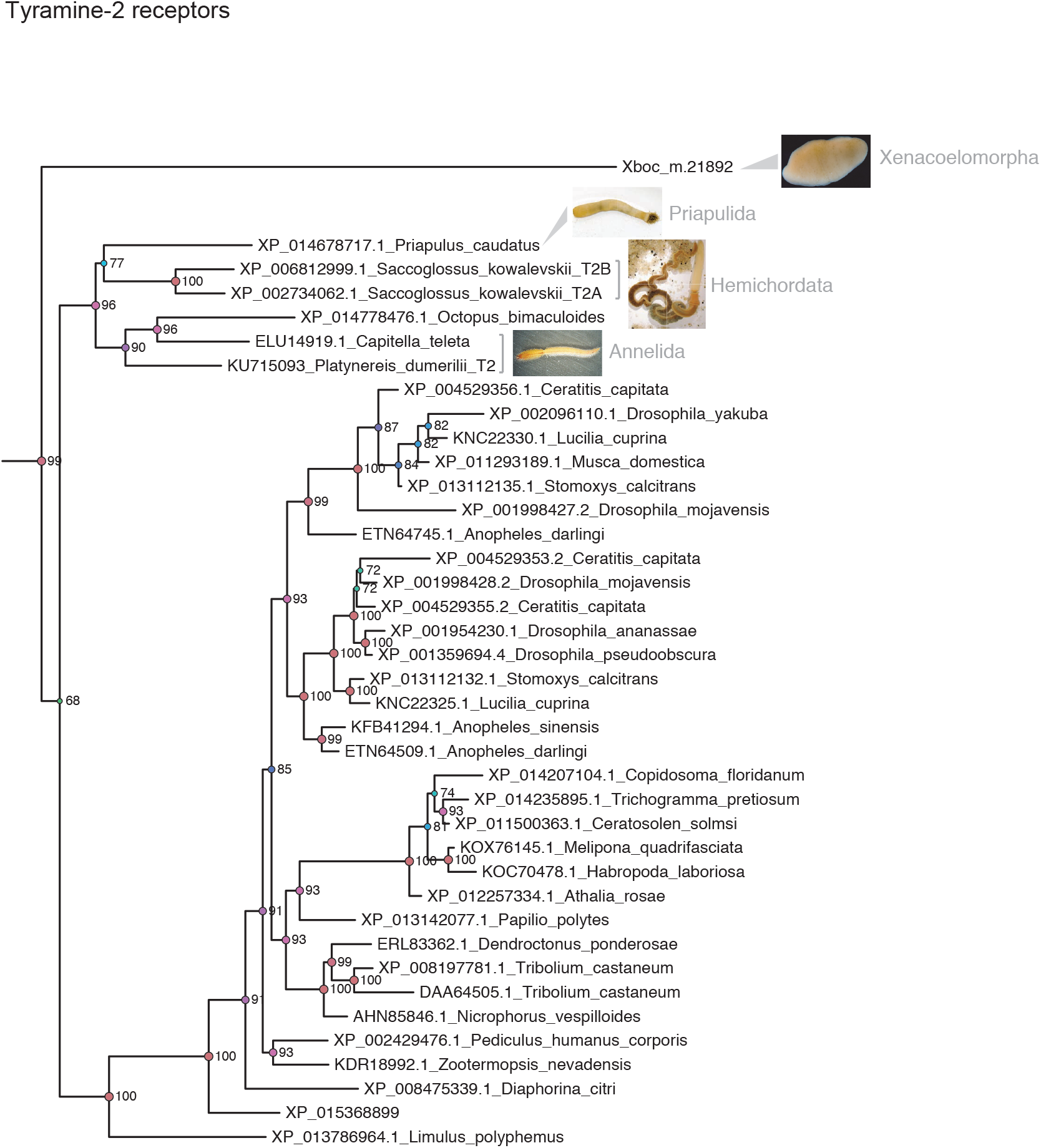
Maximum likelihood tree of Tyramine-2 receptors. Bootstrap support values are shown for selected nodes. This tree is part of a larger tree containing all investigated GPCRs. The identifiers of deorphanized tyramine receptors were tagged with T2.

**Additional file 7.**
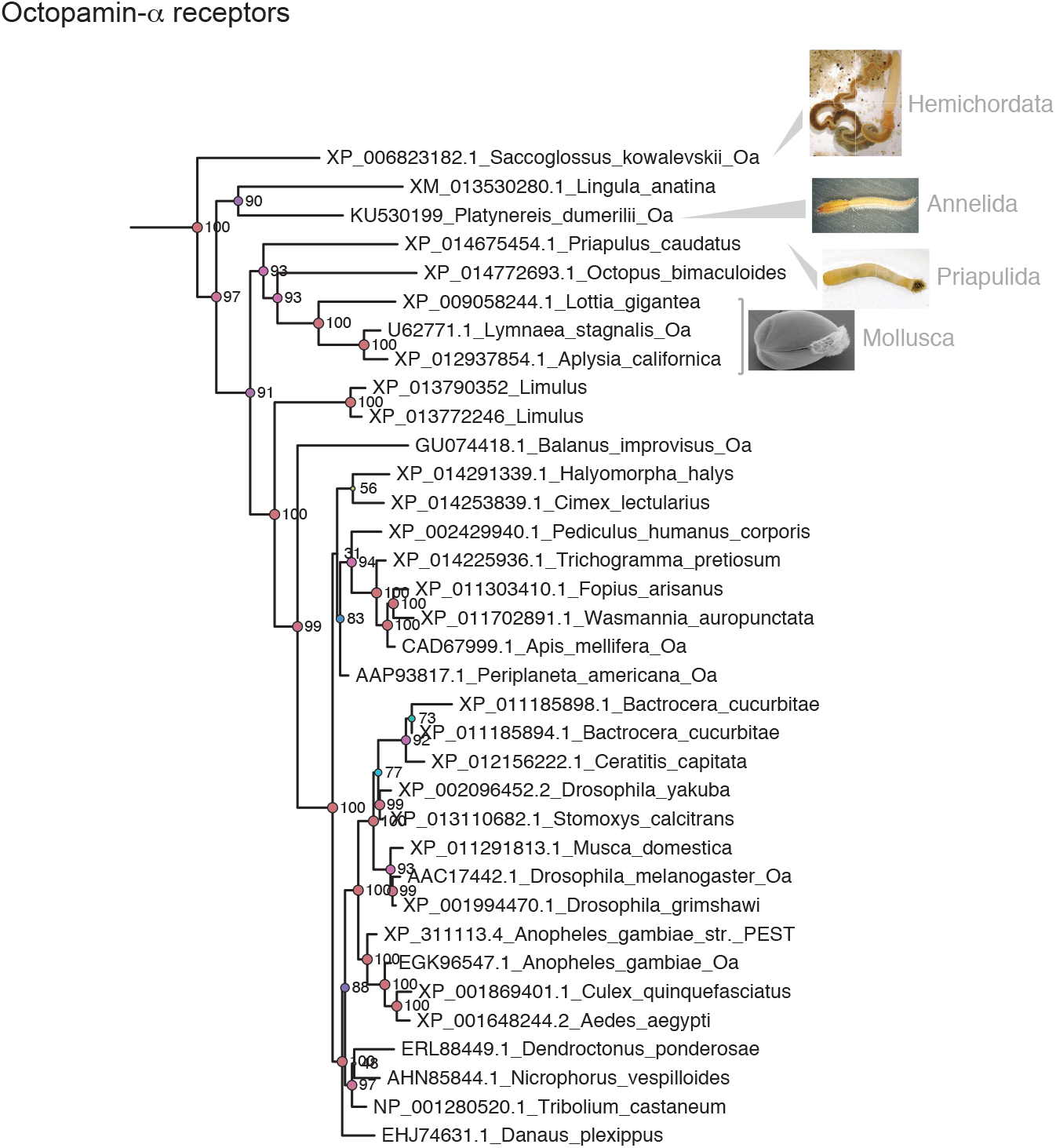
Maximum likelihood tree of Octopamine-α receptors. Bootstrap support values are shown for selected nodes. This tree is part of a larger tree containing all investigated GPCRs. The identifiers of deorphanized octopamine receptors were tagged with _Oa.

**Additional file 8.**
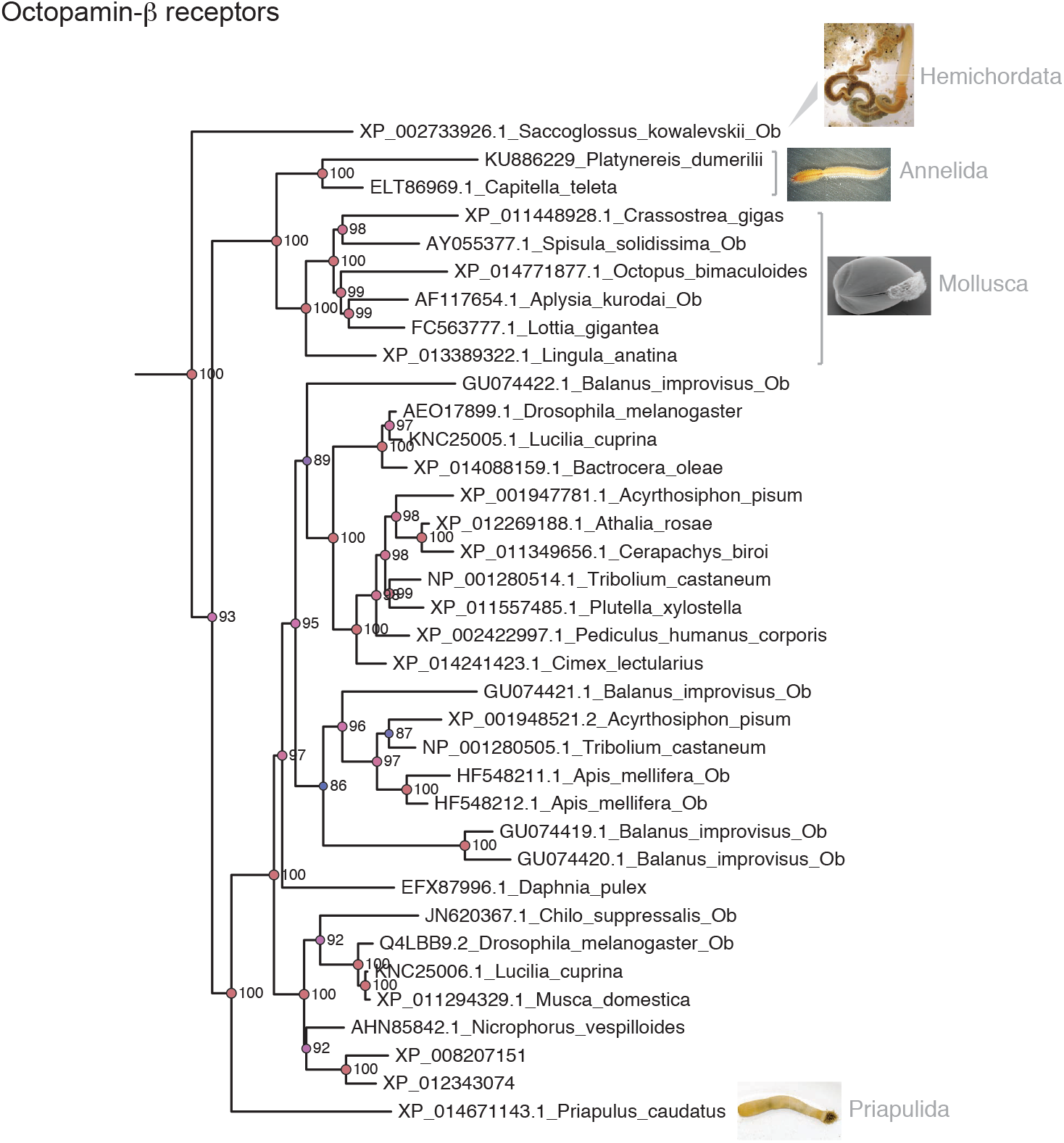
Maximum likelihood tree of Octopamine-β receptors. Bootstrap support values are shown for selected nodes. This tree is part of a larger tree containing all investigated GPCRs. The identifiers of deorphanized octopamine receptors were tagged with Ob.

**Additional file 9.**
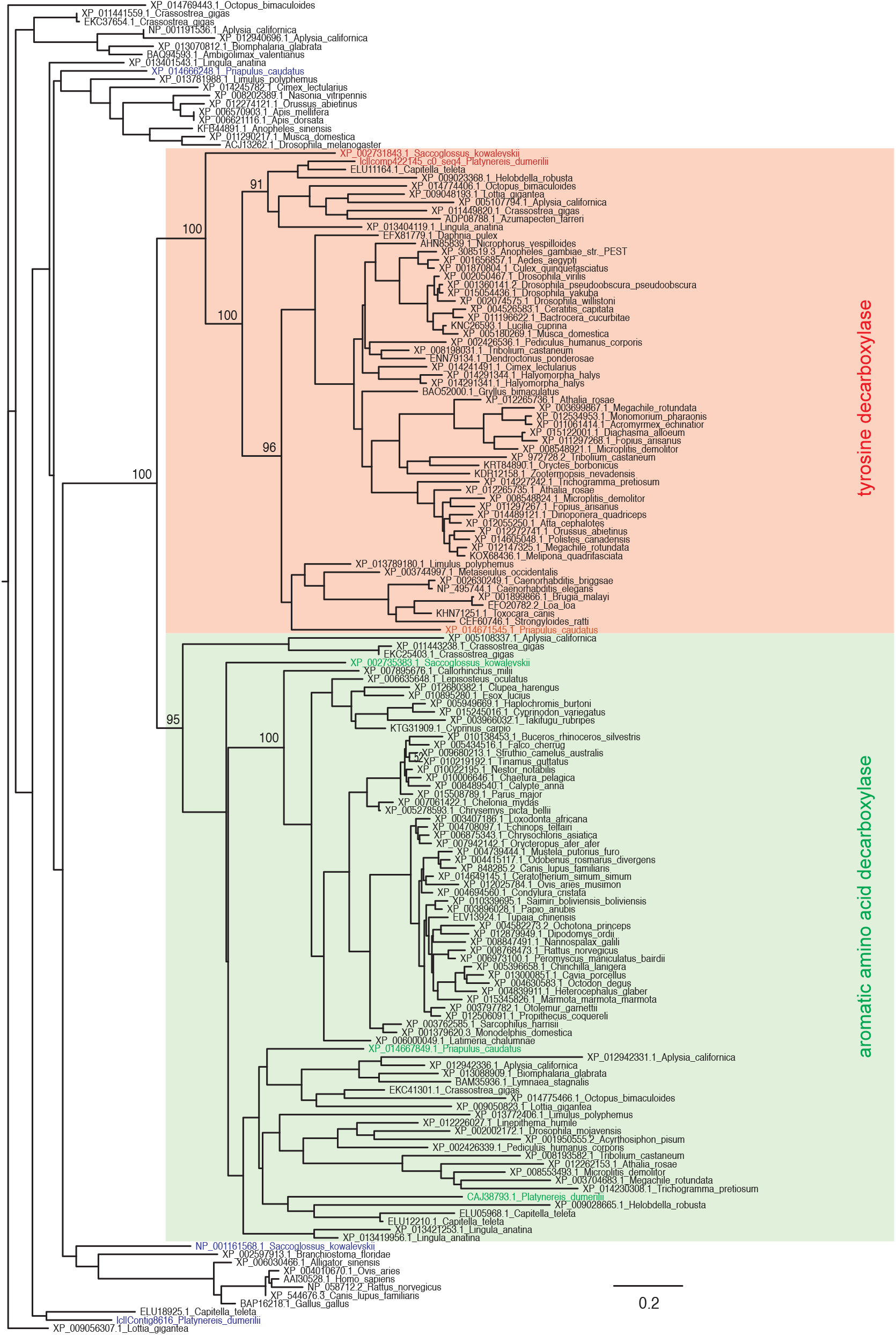
Maximum likelihood tree of tyrosine decarboxylase and aromatic amino acid decarboxylase enzymes. Bootstrap support values are shown for selected nodes. *P. dumerilii. P. caudatus*, and *S. kowalevskii* sequences are highlighted in color. The *Caenorhabditis elegans* tyrosine decarboxylase was experimentally shown to be required for tyramine biosynthesis [32].

**Additional file 10.**
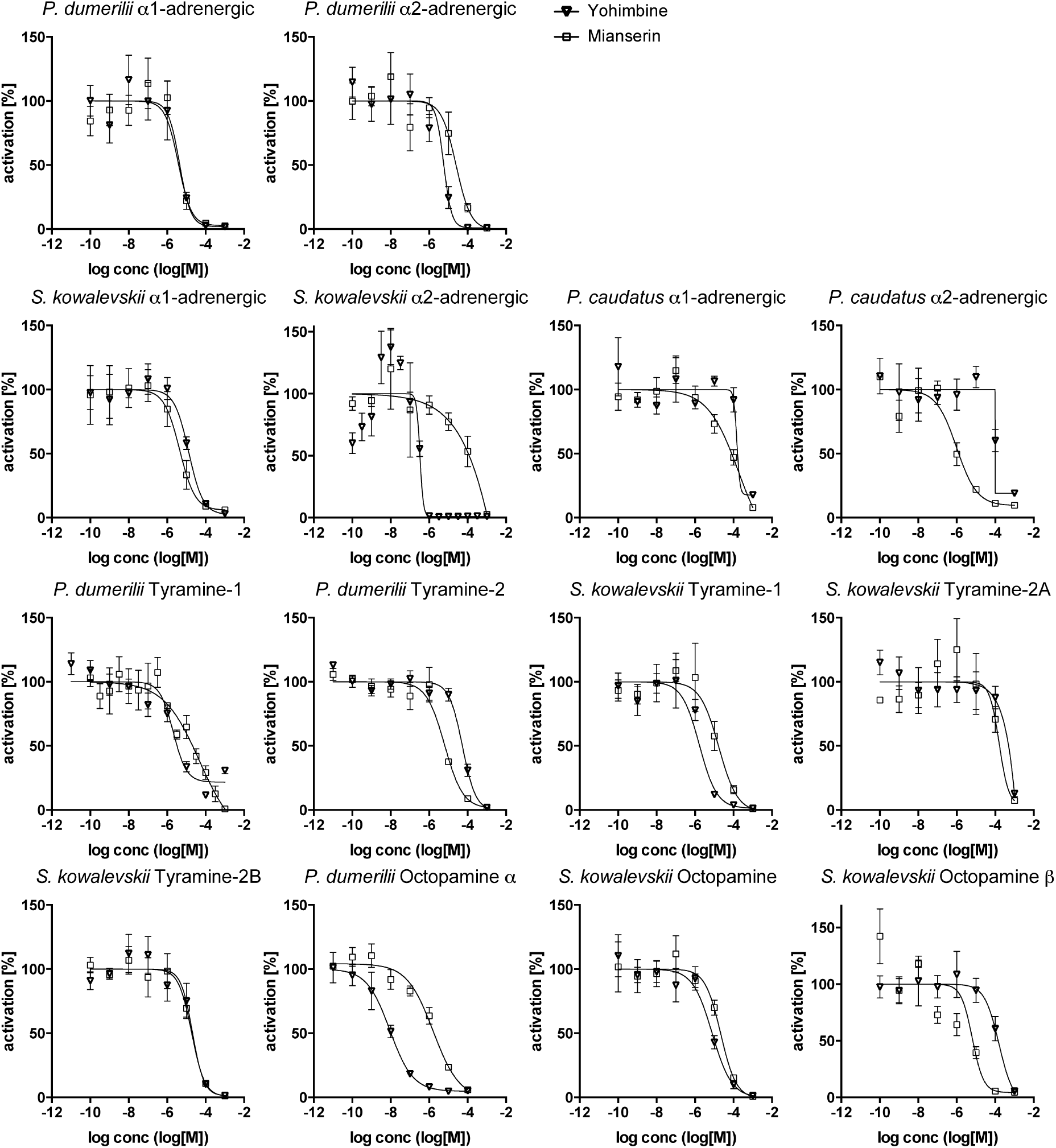
Dose-response curves of adrenergic, tyramine, and octopamine receptors from *P. dumerilii, P. caudatus*, and *S. kowalevskii* treated with varying concentrations of inhibitors. Data, representing luminescence units relative to the maximum of the fitted dose-response curves, are shown as mean ± SEM (n = 3). IC50 values are listed in Table 1.

